# Soybean Yellow Stripe-like7 is a symbiosome membrane peptide transporter essential for nitrogen fixation

**DOI:** 10.1101/2020.03.27.011973

**Authors:** A Gavrin, PC Loughlin, EM Brear, OW Griffith, F Bedon, M Suter Grotemeyer, V Escudero, M Reguera, Y Qu, SN Mohd-Noor, C Chen, MB Osorio, D Rentsch, M González-Guerrero, DA Day, PMC Smith

**Affiliations:** School of Life Sciences. La Trobe University, Bundoora, Victoria 3083. Australia; School of Life and Environmental Science, The University of Sydney, Sydney, New South Wales, 2006, Australia; College of Science and Engineering, Flinders University, Bedford Park, Adelaide, SA, Australia; IPS, Molecular Plant Physiology, University of Bern, Altenbergrain 21, 3013 Bern Switzerland; Centro de Biotecnología y Genómica de Plantas (UPM-INIA). Universidad Politécnica de Madrid. Campus de Montegancedo. Crta. M-40 km 38. 28223 Pozuelo de Alarcón (Madrid), Spain; School of BioSciences, University of Melbourne, Parkville, VIC, Australia; Department of Biological Sciences, Macquarie University, Macquarie Park, NSW, 2109

## Abstract

Legumes form a symbiosis with rhizobia that convert atmospheric nitrogen (N_2_) to ammonia which they provide to the plant in return for a carbon and nutrient supply. Nodules, developed as part of the symbiosis, harbor rhizobia which are enclosed in the plant-derived symbiosome membrane (SM), to form a symbiosome. In the mature nodule all exchanges between the symbionts occur across the SM. Here we characterize GmYSL7, a member of Yellow stripe-like family which is localized to the SM in soybean nodules. It is expressed specifically in nodule infected cells with expression peaking soon after nitrogenase becomes active. Although most members of the family transport metal complexed with phytosiderophores, GmYSL7 does not. It transports oligopeptides of between four and 12 amino acids. Silencing of GmYSL7 reduces nitrogenase activity and blocks development when symbiosomes contain a single bacteroid. RNAseq of nodules in which GmYSL7 is silenced suggests that the plant initiates a defense response against the rhizobia. There is some evidence that metal transport in the nodules is dysregulated, with upregulation of genes encoding ferritin and vacuolar iron transporter family and downregulation of a gene encoding nicotianamine synthase. However, it is not clear whether the changes are a result of the reduction of nitrogen fixation and the requirement to store excess iron or an indication of a role of GmYSL7 in regulation of metal transport in the nodules. Further work to identify the physiological substrate for GmYSL7 will allow clarification of this role.

**One sentence summary:** GmYSL7 is a symbiosome membrane peptide transporter that is essential for symbiotic nitrogen fixation that when silenced blocks symbiosome development.

## INTRODUCTION

Legumes form a symbiosis with soil bacteria, rhizobia, that allows them to access N_2_ from the atmosphere. This symbiosis is an important contributor to the biological nitrogen cycle. The rhizobia fix N_2_ via the enzyme nitrogenase to produce ammonia and provide it to the plant in return for reduced carbon generated via photosynthesis. This biological N_2_-fixation provides a large proportion of the nitrogen in the natural environment (Fowler et al., 2013) and is an important component of sustainable agricultural systems, reducing the requirement for expensive nitrogen fertilizers and the pollution that can arise from their overuse (Vance, 2001).

The establishment of this symbiosis involves signaling between the two partners and results in rhizobia moving through an infection thread derived from an invaginated root cell wall into the root cortex where a new organ, the nodule is initiated. The cell wall of the root cells is degraded and the rhizobia released into the cell. Within the nodule infected cells, the rhizobia are enclosed in a plant-derived membrane to form an organelle-like compartment called the symbiosome. Within this symbiosome the rhizobia differentiate into their symbiotic form, the bacteroid. The symbiosome membrane (SM), initially derived from the plasma membrane (PM), becomes specialized as an interface between the bacteroid and its plant host, segregating the bacteroids from the plant cytoplasm and “protecting” them from any plant defense response (Mohd-Noor et al., 2015).

The major metabolite exchange across the SM is fixed nitrogen (principally ammonia) to the plant and a carbon source, most likely malate, to the bacteroids. However, transport of many other compounds into the symbiosome across the SM must occur as the enclosed bacteroids depend on the plant for all of their nutrients, including iron, zinc, calcium and cobalt amongst others (Brear et al. 2013; Udvardi and Poole, 2013; Clarke et al., 2014). The SM effectively controls the symbiosis via a suite of transport proteins synthesized by the plant. The plant can control what moves into the symbiosome and, presumably, can withhold sustenance if required. It has been suggested that the plant can impose sanctions on non-fixing rhizobia (Kiers et al., 2003) and controlling transport across the symbiosome membrane could regulate this. It is also probable that compounds other than ammonia/ammonium move from the bacteroids to the plant (Udvardi and Poole 2013).

Transport studies with isolated symbiosomes have demonstrated the presence of a malate transporter and an ammonium channel on the SM, as well as metal ion transporters, but their molecular identity remains elusive (Udvardi and Day, 1997; Udvardi and Poole 2013; González-Guerrero et al. 2016). A number of proteomic analyses of the SM have been reported (Wienkoop and Saalbach, 2003, Catalano et al. 2004, Clarke et al. 2015) and although the earlier studies were limited by the lack of genome sequences for the legumes studied, an array of putative transport proteins have been identified. An example is LjSST1, a sulphate transporter later shown to be essential for nitrogen fixation in the *Lotus japonicus*-rhizobia interaction (Krussell et al. 2005; Schneider et al. 2019).

The most recent analysis of the soybean SM proteome (Clarke et al. 2015) identified a protein from the Yellow Stripe-like (YSL) family, Glyma.11G203400 (known then as Glyma11g31870). YSL proteins are members of the wider oligopeptide transporter family, generally considered to transport metals chelated to phytosiderophores (PS), such as deoxymugineic acid and nicotianamine (NA) (Curie et al., 2009). In monocots, PS excreted to the rhizosphere chelate ferric iron and the complexes are transported into the plant cytoplasm by YSL transporters. Maize mutants for the first characterized member of this family show a phenotype of interveinal chlorosis that is characteristic of iron deficiency and it is this phenotype that gave rise to the name Yellow-stripe 1 (YS1). YSL proteins, often localized in xylem parenchyma, can also transport other metal chelates (Dai et al. 2018; Chu et al., 2010; Sasaki et al., 2011, Zheng et al. 2012) and are important for intracellular iron transport and iron homeostasis, with both ferric and ferrous-PS complexes transported (Lubkowitz, 2011). YSL proteins are also involved in mobilization of intracellular stores of metals (Divol et al. 2013, Conte et al. 2013). YSL transporters operate through proton co-transport driven by the membrane potential (Schaaf et al. 2004) and whether localized to the PM or internal membranes, transport is always into the cell cytosol (Lubkowitz 2011).

Despite the biochemical characterization of some members of the YSL family, the functional role of other members is less clear. In particular, members of one phylogenetic clade of the YSL family, including YSL5, 7 and 8 (Group III) are not well characterized. A recent study showed that Arabidopsis YSL7 and 8 are responsible for the import of a *Pseudomonas syringae* virulence factor, syringolin A (Syl A), into the plant cytoplasm where it inhibits the proteasome (Hofstetter et al. 2013). Syl A is a peptide derivative and peptides of 4-8 amino acids in length were able to inhibit its transport in plants and in yeast expressing AtYSL7. Consequently, it was suggested that AtYSL7 and AtYSL8 act as oligopeptide transporters, although direct evidence of oligopeptide transport was not shown (Hofstetter et al. 2013).

In this study we show that both AtYSL7 and GmYSL7 (encoded by Glyma.11G203400) can transport oligopeptides and that the soybean protein, which is localized to the SM in nodule infected cells, is essential for nitrogen fixation.

## RESULTS

### GmYSL7 is a transporter of the YSL family

In our proteomic study (Clarke et al., 2015) we identified Glyma.11G203400 on the SM of soybean nodules. The protein is a member of the oligopeptide transporter (OPT) superfamily (Saier, 2000; Yen et al., 2001; Stacey et al., 2008) and has significant homology with members of the YSL family (Curie et al. 2009). We named it YSL7, as its closest Arabidopsis homologue is AtYSL7 (74% amino acid identity and 85% similarity; Yorden et al. 2011).

GmYSL7 belongs to a family consisting of 15 members in soybean (Supplementary Fig. S1; Schmutz et al., 2010) which in phylogenetic analysis fall into the three clades with both monocots and dicots members (Groups I – III, Supplementary Fig. S1). Group IV has only monocot members. In Genbank six proteins are annotated as “probable metal-nicotianamine transporter YSL7” but in the phylogenetic analysis only GmYSL7, Glyma.11G203400, associates closely with AtYSL7 in Group III, also clustering with the chickpea protein CaYSL7 (Ca08876) and three *M. truncatula* proteins (Medtr3g063490 [MtYSL7], Medtr3g063520 [MtYSL9] and Medtr5g091600 [MtYSL8]) (Supplementary Fig. S1). Of the other soybean proteins annotated as YSL7, Glyma.09G164500 and Glyma.16G212900 are more closely related to AtYSL5 and AtYSL8, while Glyma.09G281500, Glyma.20G004200 and Glyma.20G004300, although part of Group III, form a sub-clade not associated with any YSL proteins from other plants included in the phylogeny (Supplementary Fig. S1).

### GmYSL7 is expressed in infected cells of soybean root nodules

Publicly available transcriptomic data for soybean suggests nodule-specific expression of *GmYSL7* (Severin et al. 2010; Supplementary Fig. S2). We confirmed this by measuring *GmYSL7* transcript abundance in leaves, roots of 8-day old seedlings, nodules and denodulated roots of 32-day-old plants using quantitative reverse transcription (RT-q) PCR. *GmYSL7* transcript was abundant in nodules but almost undetectable in other plant organs examined (Fig 1A). We investigated the expression patterns of other YSL genes and all had lower nodule expression than *YSL7* and transcripts present in other tissues (Supplementary Fig. 2).

**Figure 1.**
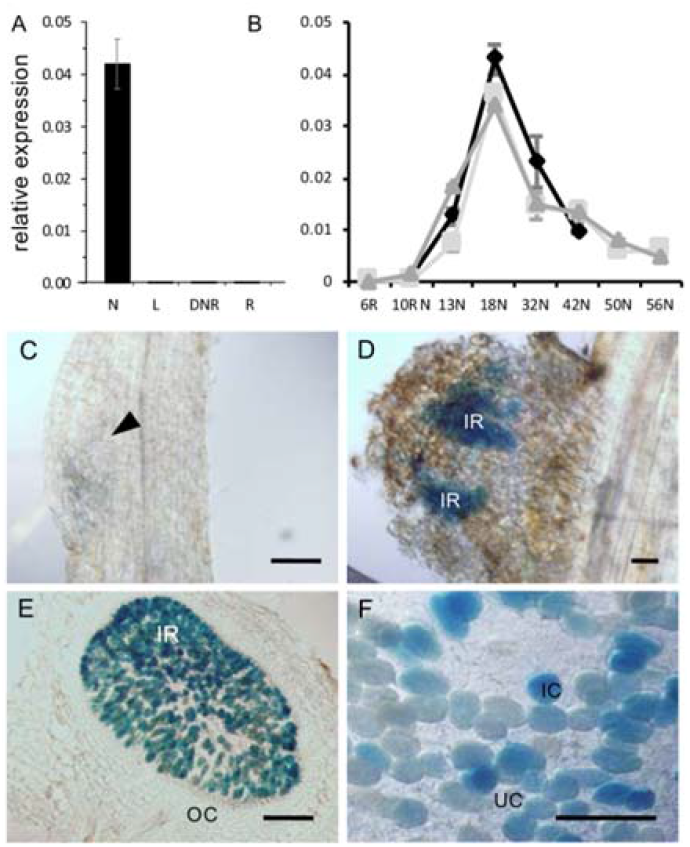
GmYSL7 is expressed in infected cells of soybean root nodules. A. Transcript level of *GmYSL7* in tissue samples from different organs. N, nodules; L, leaves; DNR, denodulated roots; R, roots. B. Transcript level of GmYSL7 during nodule development. 6R, roots 6 days after inoculation (DAI); 10RN, roots and nodules 10 DAI; 13N-26N, nodules the indicated DAI. Data shown are for three independent time courses. Bars, SE (n = 3). Nitrogenase activity was first detected at 18 DAI. C. Transgenic root expressing pYSL7:GFP-GUS. GUS staining was not detectable in the very early stages of nodule development. Arrowhead indicates a nodule initiation. D. Transgenic pYSL7:GFP-GUS 10-day-old nodule primordia. IR, infected region. E. Transgenic pYSL7:GFP-GUS mature nodule. GUS staining is restricted to infected cells. OC, outer cortex. F. Magnification of E. Scale bars, 150 μm.

Expression of *GmYSL7* during nodule development was examined with mRNA essentially undetectable in young (6 – 10-day-old) inoculated roots; however, transcript abundance increased sharply before nitrogenase activity was first detected (day 18; Supplementary Fig. 3), peaking in nodules from 18-day-old plants, and steadily decreased after this time (Fig. 1B).

As some characterized YSL proteins are involved in transport of iron complexes we examined expression of *GmYSL7* in nodules grown under varied (0 – 100 μM) added iron conditions. Two replicates were completed and although the level of expression was slightly lower in the second (not shown), the pattern of expression was similar. *GmYSL7* expression was largely insensitive to iron concentration (Supplementary Fig. 3). This was in contrast to an *AtYSL3* homologue with clear upregulation in high iron conditions (results not shown).

We investigated *GmYSL7* cellular expression pattern and the subcellular localization of the expressed protein in nitrogen-fixing nodules. The 2 kb genomic fragment immediately upstream of the coding region of GmYSL7 was inserted upstream of a promoter-less green fluorescent protein-*β*-glucuronidase (GFP-GUS) coding region to give *pGmYSL7:GFP-GUS*. GUS staining of *pGmYSL7:GFP-GUS* transformed roots and nodules agreed well with our RT-qPCR data, with no staining detectable in roots or in early nodule initials (Fig. 1C). GUS staining became evident as nodules developed (Fig. 1D) and was strongest in maturing nodules. In mature nodules, GUS staining was detected in the infected region and appeared to be confined to rhizobia-infected cells (Fig. 1E and F). No GUS staining was detected in the outer cortex of the nodule (Fig. 1E) or in untransformed nodules (data not shown).

Localization of GmYSL7 on the SM in rhizobia-infected cells was confirmed using transgenic nodules expressing *pGmLBc3:GFP-GmYSL7* and analysed by confocal microscopy. FM4-64, a lipophilic dye that fluoresces when bound to membrane (Vida and Emr, 1995), was used to counterstain the SM (Limpens et al. 2009, Gavrin et al. 2014). GFP-GmYSL7 signal was on internal membranes within infected cells but not on the PM (Fig. 2B, D). Co-localization of GFP and FM4-64 (Fig. 2A-C) signals indicates discrete localization of GFP-GmYSL7 on the SM as can be seen also in Fig 2D, with GFP-YSL forming a clear “halo” around the perimeter of symbiosomes. The GFP-YSL7 fluorescence pattern in infected cells (Fig 2D) was distinct from free GFP, detected in the cytoplasm (Fig. 2E), and from that of a construct targeted to the symbiosome space (Fig. 2F).

**Figure 2.**
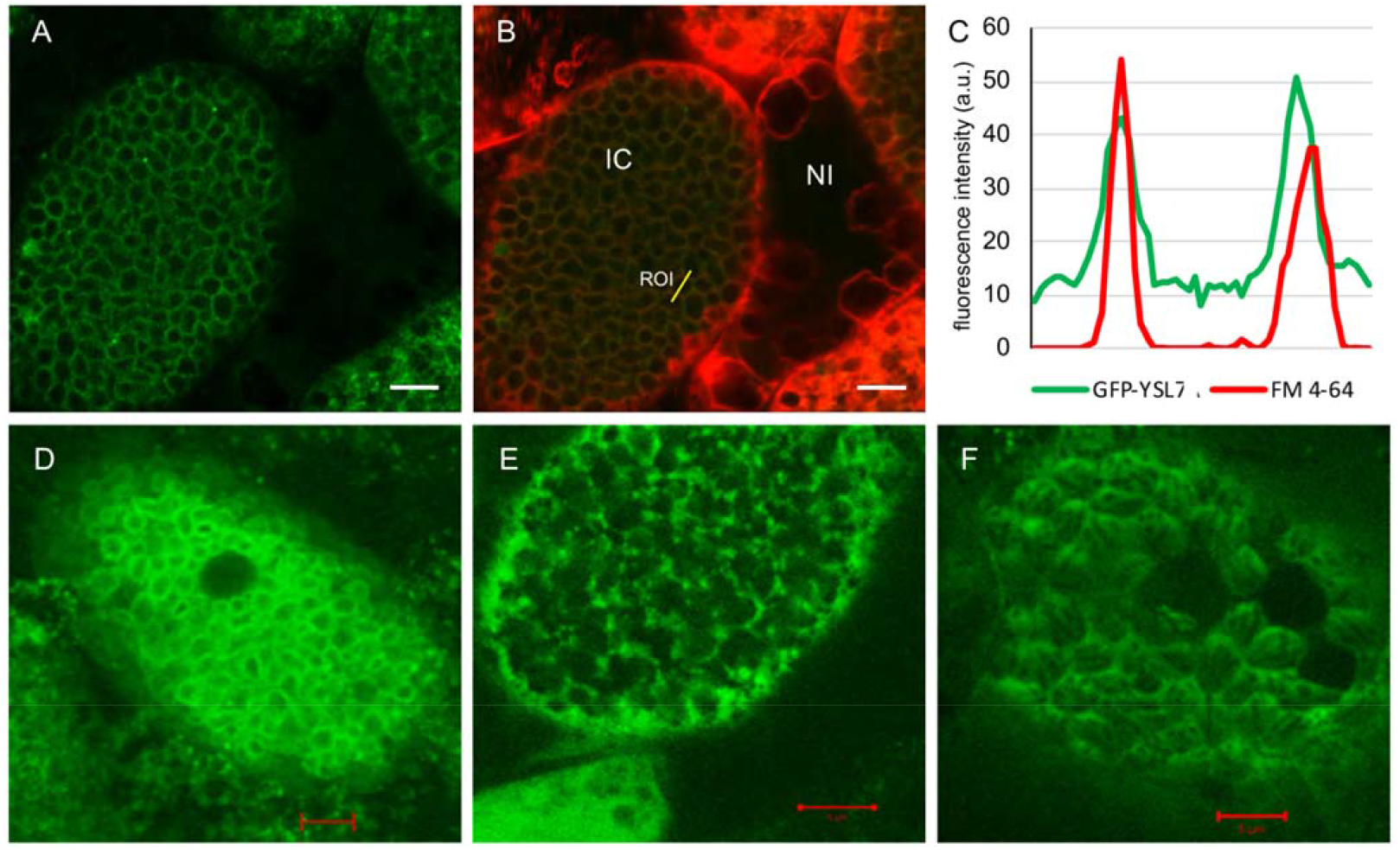
Localization of GmYSL7 in soybean nodule cells infected with rhizobia. A. GFP-GmYSL7 localizes on symbiosome membranes in infected cell of soybean nodules. B. Colocalization of GFP-YSL7 with membrane lipophilic dye FM4-64 in the same cell. IC, infected cell; NI, non-infected cell; ROI, region of interest. C. Fluorescent intensity plot of ROI from B. D. Superimposed confocal image of GFP-GmYSL7 signal on the symbiosome membrane in rhizobia-infected nodule cells. E. Free GFP localizes to the cytoplasmic spaces surrounding symbiosomes in infected cells. F. MtNOD25-GFP (Hohnjec et al. 2009) localizes to the peribacteroid space inside the symbisomes. Scale bars, 5μm.

Further confirmation that GmYSL7 was localized on the SM and not the PM was obtained by proteomic analysis of isolated SM and microsomal extract enriched in PM and endoplasmic reticulum. Approximately six times more peptides from the well characterized, SM-localized GmNOD26 were in the SM sample compared to the microsomal membrane sample, indicating enrichment of the SM in the purified sample. GmYSL7 peptides were only in the purified SM sample (Supplementary Table 1).

### Silencing of GmYSL7 interrupts development of the symbiosis

Since GmYSL7 is localized to the SM, we investigated whether it is essential for development of the symbiosis and nitrogen fixation by rhizobia using RNA interference (RNAi). Nodules from transgenic roots silenced for GmYSL7 were analyzed 24 days post inoculation (dpi; Fig. 3C). Expression of *GmYSL7* in RNAi nodules was approximately 40% of the control but expression of the closest homologs, *Glyma.16G212900 (GmYSL8)* and *Glyma.09G164500 (GmYSL5)*, was not affected (Fig. 3C). Acetylene-reduction analyses showed that nitrogenase activity was reduced in silenced nodules to only 25% of the activity of control nodules (Figure 3D). *GmYSL7* silenced nodules were smaller (Fig. 3E) and displayed a delay in development (Fig. 3A, B) in comparison to empty vector control nodules (Fig. 3 G-H).

**Figure 3.**
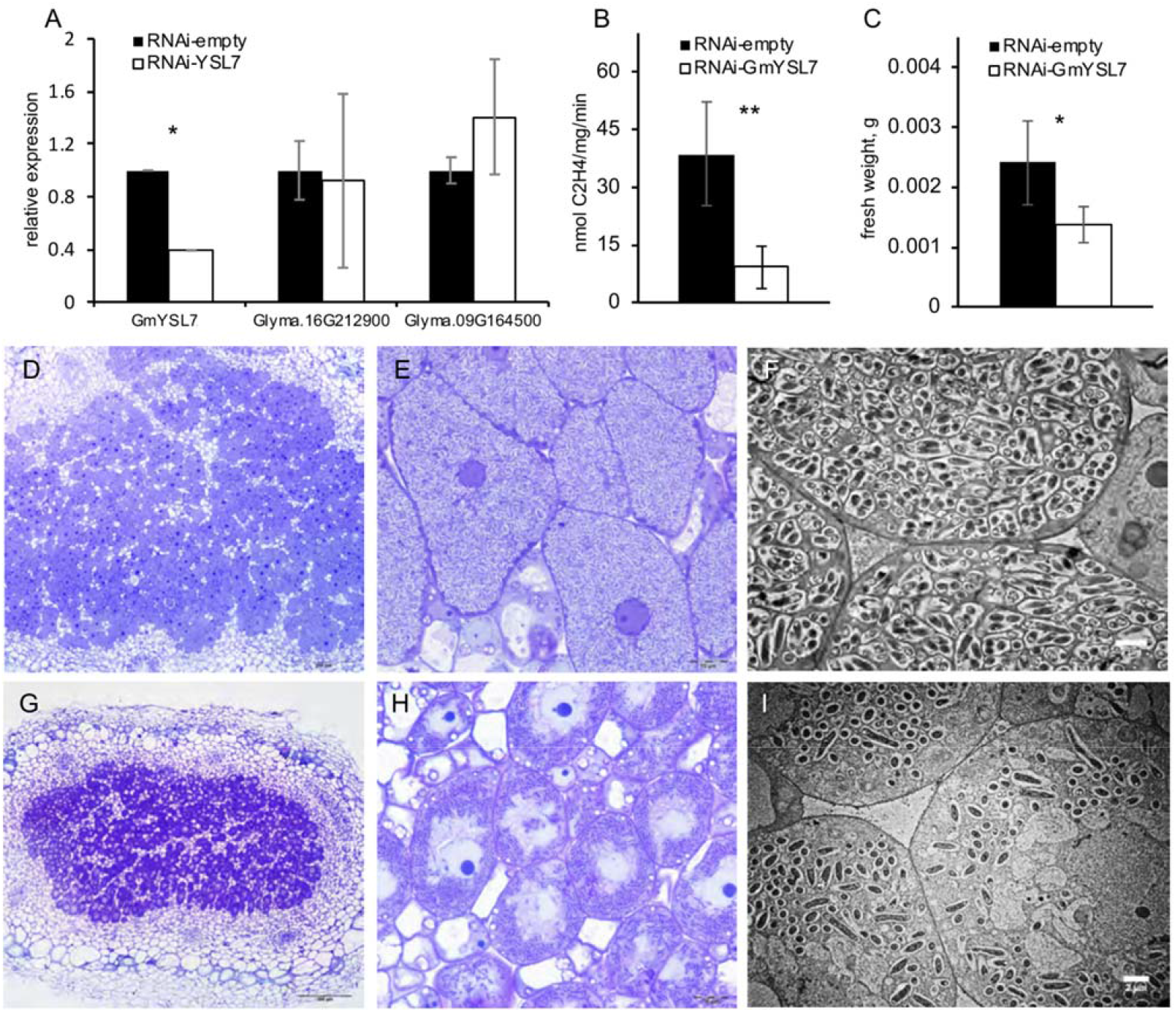
RNAi silencing of GmYSL7 affects nodule development. A. Transcript level of GmYSL7 and its closest homologs in 24-day-old nodules of empty vector control and RNAi-GmYSL7 plants (error bars represent SD; n=4; t-test: *, p<0.05). B. Nitrogenase enzyme activity in 24-day-old nodules of empty vector control and RNAi-GmYSL7 plants (error bars represent SD; n=8; t-test: **, p<0.01). C. Fresh weight of 24-day-old nodules of empty vector control and RNAi-GmYSL7 (error bars represent SD; t-test: *, p<0.05). D. Longitudinal section of a 24-day-old nodule from an empty vector control plant. E. Magnification of (A) showing developed (stage IV) infected cells. F. Electron microscopy of infected cells of a 24-day-old nodule from empty vector control containing developed multibacteroid symbiosomes. G. Longitudinal section of a RNAi-GmYSL7 24-day-old nodule. H. Magnification of (D) showing undeveloped (stage II) infected cells. I. Electron microscopy of infected cells of a RNAi-GmYSL7 24-day-old nodule containing undeveloped single-bacteroid symbiosomes. Scale bars as indicated.

Silencing of *GmYSL7* did not affect bacteria release, but infected cells remained small and, unlike the control nodules, contained small, single-bacteroid symbiosomes (Fig. 3B). Numerous small vacuoles were localized around the nucleus (Fig. 3F) whereas control nodules had no vacuoles (Fig. 3H). To pinpoint the developmental stage in wild type nodules that matches the RNAi nodules we completed an analysis of nodules from soybean infected with *Bradyrhizobium diazoefficiens* strain 1042-45 carrying the *lacZ* fusion driven by the *nifD* promoter (Acuña et al., 1987). Four stages of development were identified and images can be seen in Supplementary Fig. 4. The morphology of infected cells of *GmYSL7*-silenced nodules (Fig. 3B) appeared to be arrested at stage II of normal nodule development (Fig. 3F, Supplementary Fig. 4B, F) where numerous small vacuoles were present in infected cells and most symbiosomes contained single elongated bacteroids. Electron microscopy (EM) of the silenced nodules confirmed that the infected cells were small and under-developed, packed with symbiosomes containing only a single bacteroid (Fig. 3). Bacteroids appeared elongated as seen in control nodules during stage II (Supplementary Fig. 4F). Infected cells also contained numerous vacuoles of different sizes and apparent endosomes fusing with symbiosomes, reminiscent of the formation of a lytic compartment, which usually occurs during nodule senescence (Fig. 3I). Symbiosomes were isolated from silenced and control nodules, and this showed that in the silenced nodules, symbiosomes contained only single bacteroids compared to the control symbiosomes, which had multiple bacteroids (Supplementary Fig. 4), confirming the phenotype seen by EM analysis. The results suggest that silencing of *GmYSL7* arrests development of soybean nodules at stage II.

### RNAseq of nodules in which *GmYSL7* is silenced

We used RNAseq to compare the transcriptome in 22-day old nodules from *GmYSL7*-RNAi plants and empty vector controls. In these experiments, *GmYSL7* expression in RNAi plants was around half that of control expression. Principal component analysis (Fig. 4A) and a heatmap of gene expression of all differentially expressed genes shows clear differences between the RNAi and control samples (Fig. 4B). There were no significant changes in expression of other YSL genes.

**Figure 4.**
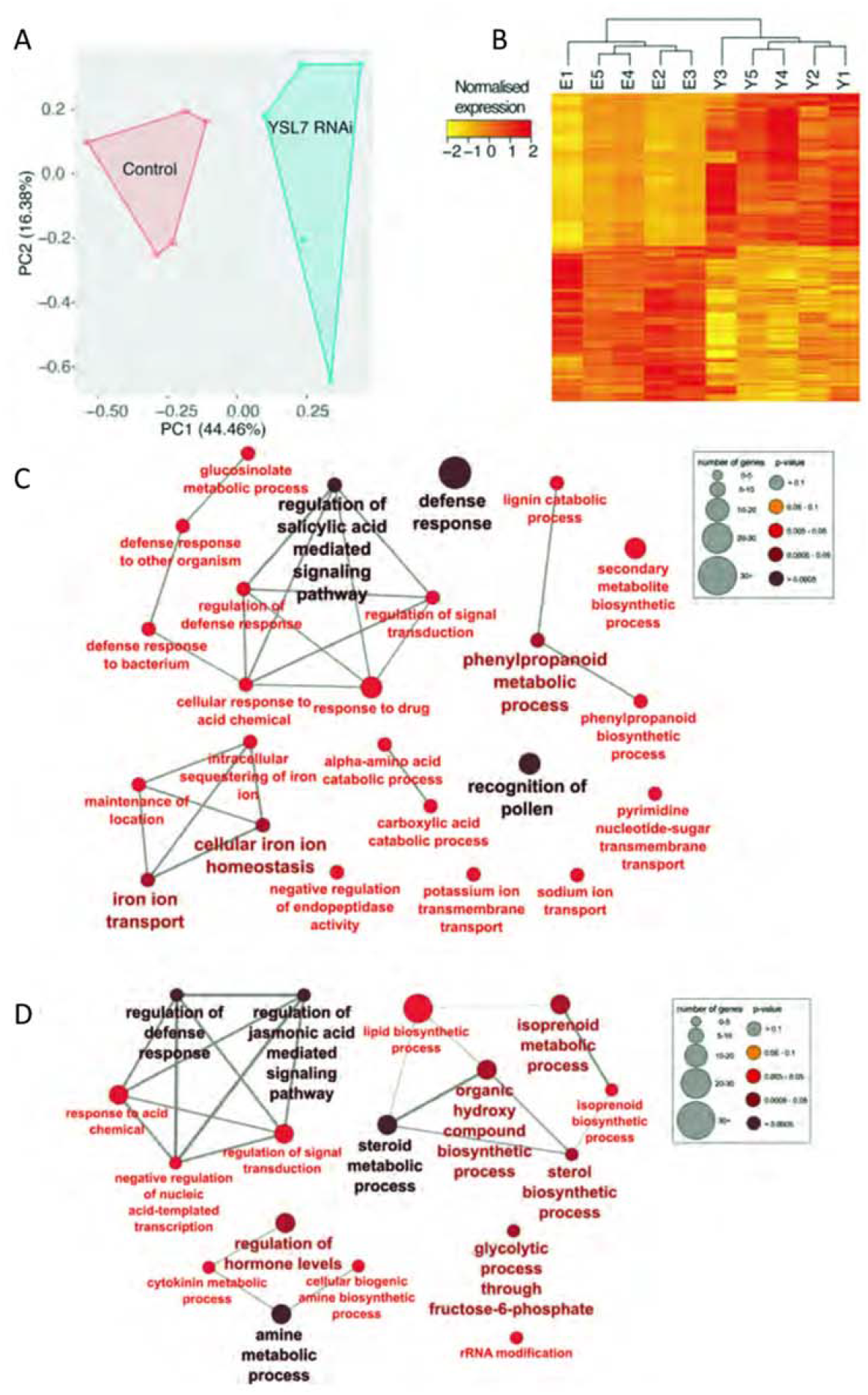
RNAseq analysis of *GmYSL7-*RNAi nodules. A. Principal component analysis of RNAseq samples. B. Heatmap of clustering of differentially regulated genes in each nodule sample. Gene-expression values are normalized by using a z-score transformation on TPM. C, D. Gene Ontology enrichment analysis of biological processes in up (C) and down (D) regulated genes from GmYSL7 RNAi nodules. GO term enrichment analyses were performed using the ClueGO v2.5.5 plugin (Bindea et al., 2009) in Cytoscape v3.5.1 (Shannon et al., 2003). Circles represent an enriched group of genes based on their GO terms. Circle size and colour indicate the number of mapped genes and associated Term PValue corrected with Bonferroni step down.

There were 924 genes with log_2_fold change of 1 or greater in nodules in which YSL7 was silenced, while 1180 genes had log_2_fold change of −1 or greater (with adjusted p-valiue <0.05). Gene ontology (GO) enrichment analysis showed that genes involved in defence responses (e.g. defence response to bacterium, defence response to other organism), “negative regulation of endopeptidase activity” and a network associated with iron homeostasis, sequestration and transport, are overrepresented in the upregulated genes (Fig. 4C). A network of genes with GO terms associated with regulation of transcription and signal transduction, and another including those associated with lipid biosynthesis, are overrepresented in the downregulated transcripts (Fig. 4D).

Details of expression of particular genes in the RNAi and control nodules are available in Supplementary Table 2. Among the genes with significantly higher expression in the silenced nodules were those encoding homologues of a senescence-associated gene 13 (Glyma.12G059200), NRT1.8/NPF7.2 (proton-coupled H+/K+ antiporter, Glyma.18G260000), organic cation/carnitine transporter4 (Glyma.12G216400), ferritin (Glyma.01G124500, Glyma.11G232600, Glyma.03G050100), vacuolar iron transporter-like proteins (Glyma.05G121200, Glyma.08G076000), plantacyanin (Glyma.08G128100), a copper transport protein (Glyma.09G179800), cation efflux family protein (Glyma.08G164800), a cationic amino acid transporter 2 (Glyma.19G116500), nitrate transporter 2.4 (Glyma.11G195200) and a number of protease inhibitors (Supplementary Table 2).

Genes with significantly lower expression in the *GmYSL7* silenced nodules were those encoding homologues of sucrose-proton symporter 2 (Glyma.16G156900), glutamine dumper 2 (Glyma.18G277600), nicotianamine synthase 1 (NAS, Glyma.15G251300), GmNIC1a (Glyma.12G208900), a Clavata3/ESR (CLE) related homologue, and Putative lysine decarboxylase family protein (Supplementary Table 2).

### GmYSL7 transports oligopeptides and Syringolin A but not Fe(II)-NA

Yeast complementation was used to try to identify a substrate for GmYSL7. Initially we tested for transport of Fe(II)-NA by complementation of the *fet3/fet4/ftr1* mutant; however, although the positive control *ZmYS1* (Curie et al. 2001, Schaaf et al. 2004) complemented the mutant, *GmYSL7* and *AtYSL7* did not (Fig. 5).

**Figure 5.**
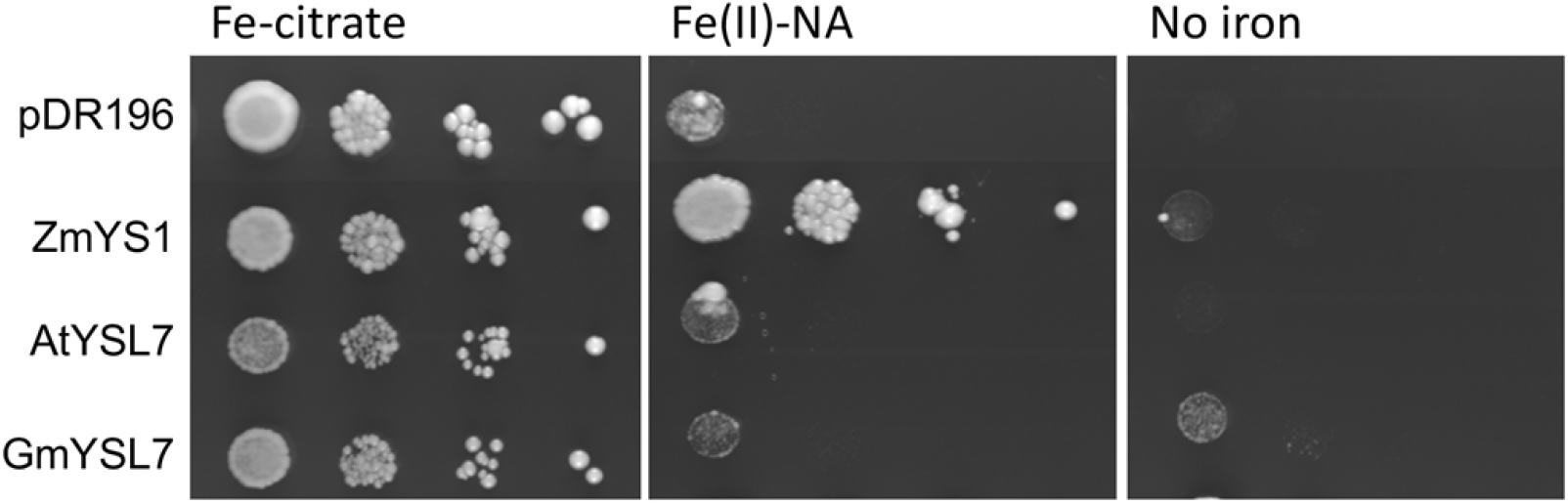
GmYSL7 does not transport iron or Fe(II)-nicotianamine (NA). DEY1530 yeast (*fet3/fet4/ftr1*) was transformed with the empty vector plasmid pDR196, *AtYSL7, GmYSL7* or *ZmYS1* in pDR196GW. Serial dilutions of each yeast transformant were applied to SD plates (that include 1.6 μM FeCl_3_) with 10 μM Fe-citrate, Fe(II)-NA or no added iron (no iron) and the plates grown for 3-5 days.

Since the YSL family is part of the wider oligopeptide transporter (OPT) family, we next tested whether GmYSL7 could complement the yeast oligopeptide transport *opt1* mutant, using different oligopeptides as the sole source of nitrogen for growth. When the transformants were grown with four (ALAL, LSKL), five (IIGLM) and six (KLLLLG) amino acid peptides as the only N source, cells expressing AtYSL7, GmYSL7 or AtOPT4, but not with the empty vector pDR196, grew (Fig. 6). On media containing larger peptides (eight, ten or twelve amino acid), the growth of the transformants was more varied. AtOPT4 supported growth on the eight amino acid peptide DRVYIHPF, while growth was weak for AtYSL7 and GmYSL7. Growth on the 10 amino acid peptides DRVYIHPFHL was close to background for all transformants, but all grew better than vector control on the 12 amino acid peptide RLAPEGPDPHHN (Fig 6) which corresponds to the mature CLE peptide, GmRIC1a (encoded by Glyma.13G292300; Hastwell et al. 2015).

**Figure 6.**
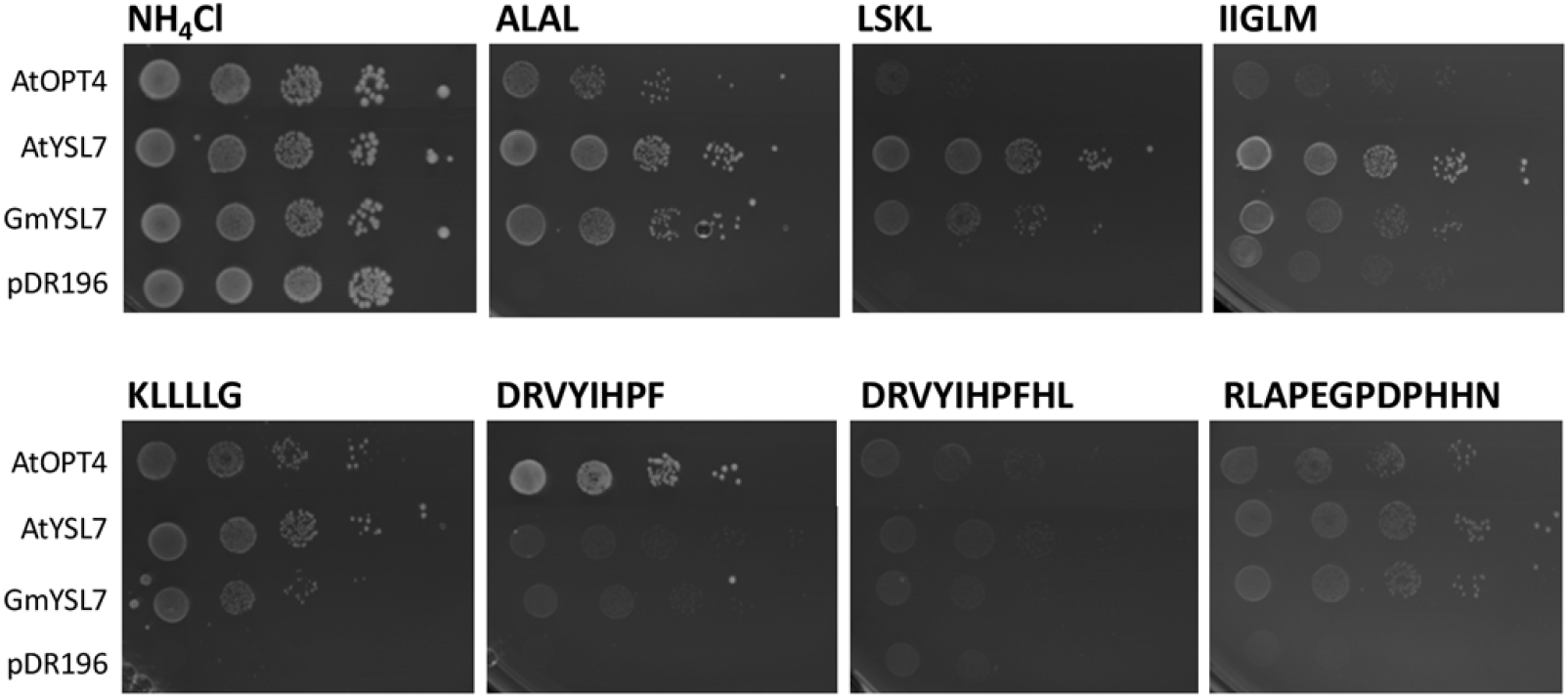
GmYSL7 and AtYSL7 transport oligopeptides. *AtOPT4, AtYSL7, GmYSL7* in pDR196GW and the empty vector (pDR196) were introduced into the yeast *opt1* mutant, Y11213. Serial dilutions of each transformant were grown as above on minimal medium containing either 10 mM NH_4_Cl (positive control) or 100 μM peptide (with sequence as indicated) as the sole source of nitrogen

AtYSL7 is involved in syringolin A (SylA) uptake and, when expressed in yeast, exposure to SylA inhibited growth (Hofstetter et al. 2013, Fig. 7) suggesting the transporter mediated uptake of this toxic peptide derivative. We used this assay to test for transport of SylA by GmYSL7. GmYSL7, AtYSL7 and the empty vector pDR195 were expressed in the yeast *pdr5* mutant, that lacks the ABC transporter PDR5, and plated as a lawn (Hofstetter et al., 2013). When a disk containing SylA was placed on the plate, growth of yeast expressing GmYSL7 and AtYSL7, but not the empty vector pDR195, was inhibited. Inhibition of growth caused by SylA on the AtYSL7 plate showed as clear patches on the plate for all concentrations of SylA tested, while inhibition of yeast expressing GmYSL7 was weaker (Fig. 7).

**Figure 7.**
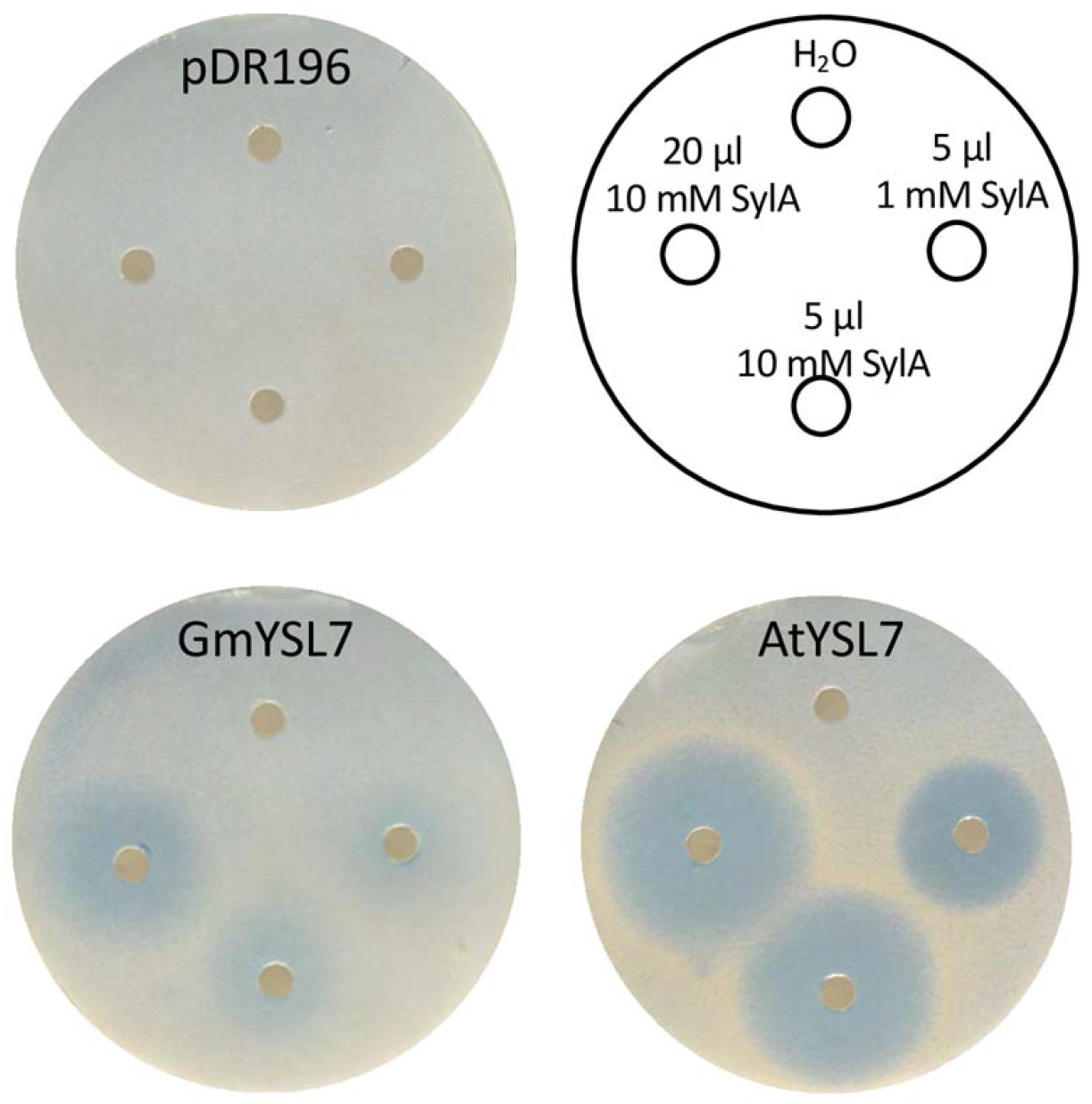
GmYSL7 transports Syringolin. A. BY4742 yeast *Δpdr5:KanMX6* transformed with the empty vector (pDR196) or the vector expressing *AtYSL7* or *GmYSL7* were plated as a lawn on solid synthetic defined (SD) media. Filter disks with the indicated SylA solutions were placed onto the plates and inhibition of growth examined after 2 days.

### GmYSL7 and MtYSL7 are functionally equivalent

In the accompanying manuscript (Castro-Rodríguez et al. 2020), MtYSL7, which is localized on the PM in the vasculature and nodule cortex in *Medicago truncatula*, is described. To determine whether GmYSL7 and MtYSL7 proteins play similar roles in the different cell types in which they are located, we expressed *GmYSL7* in the *Mtysl7* mutant. Expression was driven by the *MtYSL7* promoter to ensure expression in the cells in which MtYSL7 is present (vasculature and nodule cortex but not infected cells). Although GmYSL7 localizes to the SM in soybean, it was able to complement the *Mtysl7* transposon insertion mutant to restore nitrogenase activity to wild type levels and increase the dry weight of the transformed plants compared to the mutant (Fig. 8). Based on this result, we assume that in the *Mtysl7* mutant, when expressed in the cells where MtYSL7 is normally active, GmYSL7 at least partially localizes to the PM.

**Figure 8.**
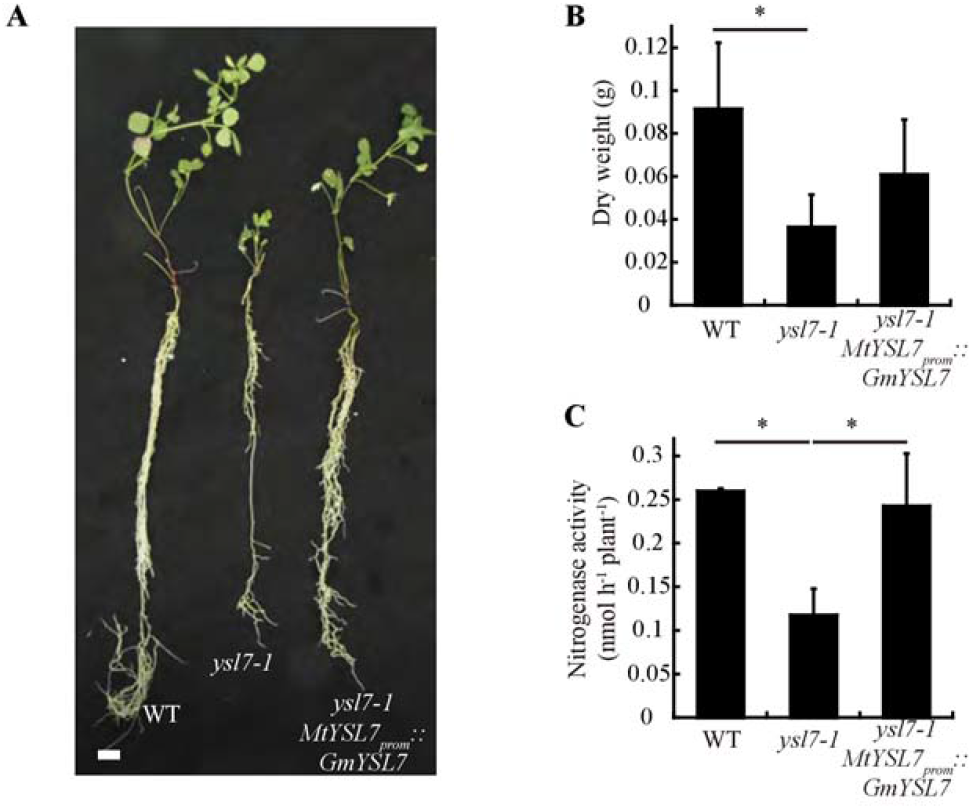
GmYSL7 is functionally equivalent to MtYSL7. A. Growth of representative wild type (WT), *ysl7-1*, and *ysl7-1* transformed with *GmYSL7* controlled by the *MtYSL7* promoter (*ysl7-1 MtYSL7_prom_∷ GmYSL7*). Bar = 1 cm. B. Dry weight of 28 dpi WT, *ysl7-1*, and *ysl7-1 MtYSL7_prom_∷ GmYSL7* plants. Data are the mean ± SE of 5 transformed plants. C. Nitrogenase activity of 28 dpi WT, *ysl7-1*, and *ysl7-1 MtYSL7_prom_∷ GmYSL7* plants. Acetylene reduction was measured in duplicate from two sets of three-four pooled plants. Data are the mean ± SE. * indicates statistically significant differences (p<0.05)

## DISCUSSION

We have characterized a member of the YSL family, GmYSL7, in soybean. The protein is part of a clade of YSL proteins (Group III) that includes AtYSL5, AtYSL7 and AtYSL8. AtYSL7 and 8 are involved in transport of the *Pseudomonas syringae* virulence factor into Arabidopsis cells across the PM (Hofstetter et al. 2013), but their physiological role in plants has not been determined. GmYSL7, AtYSL7, CaYSL7 and three *Medicago truncatula* proteins (MtYSL7, 8 and 9) form a cluster in phylogenetic analyses, but the soybean protein’s expression profile is distinct from that of *AtYSL7* and *MtYSL7*. *AtYSL7* is expressed mainly in flowers but also in siliques and roots. *MtYSL7*, has highest expression in nodules, but is also expressed in roots. Soybean, on the other hand, appears to lack a *YSL7* paralog with expression similar to *AtYSL7* and *MtYSL7*. Rather, *GmYSL7* expression is linked specifically to symbiotic nitrogen fixation, occurring only in infected nodule cells where the protein is present on the SM, but not the PM, in contrast to AtYSL7 and MtYSL7. Furthermore, its expression is only marginally affected by the iron concentration of the growth medium (Fig S4). This seems a clear example of neofunctionalization with the loss of the paralog (Xu et al. 2017). Either there is no requirement in soybean for the role played by AtYSL7 in other organs or another soybean gene with functional redundancy fulfils that role. The closest homologues of *GmYSL7*, *Glyma.16G054200* (*GmYSL8*) and *Glyma.19G094800* (*GmYSL5*) are expressed in almost all tissues (Fig. S2), but we know nothing about their function at this stage.

An important role for GmYSL7 in nitrogen-fixing nodules is shown by knockdown of its expression, which resulted in smaller nodules and a decrease in nitrogenase activity. The *GmYSL7* RNAi nodules appear to have been developmentally arrested, with small symbiosomes that contain only one bacteroid, in contrast to the large symbiosomes containing multiple bacteroids in infected cells of control nodules. The *GmYSL7* RNAi infected cell ultrastructure is similar to control nodules in the early stages of development. This suggests that the activity of GmYSL7 – and the substrate(s) it transports across the SM - is important for the continued development of the symbiosis and maturation of infected cells.

Our results show clearly that GmYSL7 transports a range of small peptides. When considering the activity of SM transporters, it is important to bear in mind the orientation and energization of the SM (Udvardi and Day, 1997), as this influences the direction that any given substrate is transported. A P-type ATPase on the SM together with the rhizobial electron transport chain, pump protons into the symbiosome space, creating an electrochemical gradient across the SM, with the membrane potential positive on the inside and the interior of the symbiosome (symbiosome space) acidic (Udvardi and Day, 1997). All YSL proteins characterised to date transport compounds across cell membranes into the cytoplasm (Lubkowitz 2011), with proton symport the most likely mechanism (Schaaf et al. 2004). Assuming that GmYSL7 has a similar mechanism, it is consequently likely to transport its peptide substrate out of the symbiosome and into the plant cell cytosol. The phenotype seen in *GmYSL7*-RNAi nodules is, therefore, related to the lack of provision of this substrate to the plant cell.

The *Medicago truncatula* homologue of GmYSL7, MtYSL7, is characterized in an accompanying manuscript and although it is not localized on the SM, the *Mtysl7*-mutant has a phenotype that also affects the symbiosis and nitrogen fixation (Castro-Rodríguez et al. 2020). The difference in cellular localization might be explained by the fact that *M. truncatula* produces indeterminant nodules, where the meristem continues to be active throughout development, and soybean determinant nodules, in which mature nodules have no meristem. As well as structural differences the different nodule types have a number of metabolic differences including in the mechanism for nitrogen assimilation and the compounds transported from the nodules (Smith and Atkins, 2002). However, *GmYSL7* is functionally equivalent to *MtYSL7* because it complemented the *Mtysl7-1* mutant, restoring nitrogenase activity and growth in low N conditions. This suggests that the two YSL7 proteins are able to transport the same substrate/s and that while MtYSL7 brings this substrate into the cell across the PM, GmYSL7 moves its substrate out of the symbiosome and into the cytosol.

The fact that *GmYSL7* complements the *Mtysl7* mutant also suggests that the mechanism for targeting the GmYSL7 protein to the PM in non-infected cells (where it is expressed from the *MtYSL7* promoter) must be modified in infected cells to allow it to reach the SM. This could involve a chaperone specific to the infected cell but might also be related to the fact that the SM is initially derived from the PM (Brear et al. 2013, Mohd-Noor et al. 2015).

In contrast to many other YSL proteins, neither of the legume YSL7 proteins, nor that from Arabidopsis, were able to transport Fe(II)-NA (this study, Castro-Rodríguez et al. 2020 accompanying manuscript). Additionally MtYSL7 could not transport Fe(III), zinc or copper complexed with NA (Castro-Rodríguez et al. 2020, accompanying manuscript). On the other hand, complementation of the yeast *opt1* mutant showed that the three proteins could transport oligopeptides of various sizes, including a CLE peptide, GmRIC1a. Like AtYSL7, when expressed in yeast, GmYSL7 could also transport syringolin A, a peptide derivative that is the virulence factor for *Pseudomonas syringae* (Hofstetter et al. 2013). Inhibition of yeast growth caused by the transported SylA was not as strong as for AtYSL7 suggesting that GmYSL7 may not transport it as effectively or have the same specificity for the compound. In our assays for direct uptake of oligopeptides in yeast, both AtYSL7 and GmYSL7 supported growth on media with oligopeptides of 4-6 and 12 amino acids as their sole N source, but there was little growth when the 8 amino acid peptide, DRVYIHPF was used, despite its ability to reduce the effect of SylA on Arabidopsis roots (Hofstetter et al., 2013). MtYSL7 was also identified as an oligopeptide transporter and showed similar specificity for oligopeptides to AtYSL7 and GmYSL7 (Castro-Rodríguez et al., 2020 accompanying manuscript).

While it is clear that YSL7 proteins are peptide transporters, their physiological role in legumes is not clear. Glutathione is a three amino acid peptide derivative found in nodules and bacteroids but as MtYSL7 cannot transport GSH (Castro-Rodríguez et al., 2020 accompanying manuscript) and GmYSL7 is able to replace the function of MtYSL7, it is unlikely that transport of GSH out of the symbiosome is the physiological role of GmYSL7. The symbiosome contains a number of proteases on the SM and in the symbiosome space (peribacteroid space; Clarke et al. 2015) and appears to act like a vacuole containing large amounts of free peptides (Clarke et al. 2015). Some of these could be substrates for GmYSL7, but why blocking their exit from the symbiosome would inhibit N-fixation and symbiosome development to such an extent is not obvious. While it is possible that GmYSL7 acts to scavenge N by transporting peptides from the symbiosome space into the plant cytosol, it is unlikely that this would have such a profound effect on nodule development.

It is tempting to speculate that release of peptides from the symbiosomes has a more direct role in manipulating plant gene expression and organogenesis. Cyclic peptides act as signaling molecules in some symbioses (Abbamondi et al. 2014) and it is possible that GmYSL7 transports an oligopeptide derivative produced in the bacteroids. In this scenario, release of the oligopeptide signal could be required to relieve plant inhibition of bacteroid division or as a positive signal for symbiosome development. Supporting this idea is the fact that a protein annotated as an oligopeptide transporter was specifically induced in symbiotic *Bradyrhizobium japonicum* (Pessi et al. 2007) and a number of transcription factors are upregulated in nodules of *GmYSL7* RNAi plants.

We used RNAseq of GmYSL7-RNAi nodules to investigate further the effects of inhibiting transport by GmYSL7. Overrepresented GO terms in the downregulated genes include a range of terms associated with lipid metabolic processes (lipid biosynthetic process, isoprenoid metabolic process). It is likely that this relates to the failure of the symbiosome to develop with multiple bacteroids. The change from a single bacteroid symbiosome to one with multiple bacteroids is likely to require synthesis of large amounts of lipid. With development of the infected cell blocked at an early stage this synthesis would not be required.

Overrepresented terms in the upregulated genes included “defense response”, “defense response to bacteria”, “defense response to other organism” and “regulation of defense response”, suggesting that blocking transport by GmYSL7 causes a general defense response against the rhizobia. This may be an indirect effect of a decrease in nitrogen fixation, with the plant sanctioning the bacteria for not doing its job (Kiers et al. 2003). Upregulation of Glyma.11G195200, a soybean homologue of AtNRT2.4, a nitrate transporter that is upregulated in response to nitrogen starvation, suggests that as nitrogenase activity was reduced, the nodules in *YSL7*-RNAi plants were indeed nitrogen starved.

In relation to oligopeptide transport, the gene encoding the CLE peptide GmNIC1a is downregulated (9.3-fold lower) in the RNAi nodules. This peptide is responsible for regulation of nodulation in response to nitrate and its expression is induced by nitrate. Over-expression of the peptide reduces nodulation (Reid et al. 2011), so lower transcript levels might be expected to favour nodule formation and development. It is interesting that YSL7 can transport GmRIC1a, a homologue of GmNIC1a. It is possible that GmRIC1a or a structurally similar peptide is synthesised in the symbiosome and exported via YSL7 to the plant cytosol where it influences nodule development. This would explain the arrested development of nodules in which YSL7 is silenced. If a GmRIC1a-like peptide is a substrate for GmYSL7, then blocking its transport may also affect expression of GmNIC1a gene. In addition to this, the GO term “negative regulation of endopeptidase activity” is overrepresented with higher expression of four protease inhibitors in the RNAi nodules. This may suggest that the peptide transported by YSL7 is processed in the plant cell and blocking its transport leads to expression of inhibitors of proteases.

Another group of GO terms that are overrepresented in the upregulated genes in GmYSL7-RNAi nodules are “intracellular sequestering of iron ion”, “iron ion transport” and “cellular iron ion homeostasis”. Some of the genes associated with these terms include ferritins and vacuolar iron transporter (VIT) gene homologues. This is accompanied by upregulation of a number proteins potentially associated with metal transport (Cu transport protein, cation efflux family protein, MATE efflux family protein) and a transcription factor, WRKY9, associated with a GO term “cellular response to iron ion starvation”. This suggests that metal homeostasis is dysregulated in the *YSL7*-RNAi nodules in a similar manner to that seen in *Mtysl7-2*, where iron and copper concentrations are increased in nodules (Castro-Rodríguez et al. 2020). There are two possible explanations for this. If symbiosome development is stalled and nitrogen fixation blocked, then metals being supplied to the symbiosome by the plant may accumulate in the nodules and need to be sequestered to avoid cellular damage. This may result in storage, particularly of iron, in uninfected cells with ferritin, or transport into the vacuole via VIT proteins.

Another intriguing effect of the silencing of YSL7 is the downregulation (16-fold decrease) of Glyma.15G251300, a gene encoding NAS, responsible for synthesis of NA, a PS involved in metal transport by YSLs. This is further evidence that YSL7 silencing affects iron homeostasis in the plant (see above). It is also possible that the downregulation in YSL7-RNAi nodules is due to changes in activity of another YSL transporter involved in metal supply to the nodule, but there do not appear to be significant changes in expression of other YSL genes in the RNAi nodules. However, it is also possible that the changes in expression of these genes are simply a consequence of the block in nodule development that occurs in nodules where YSL7 is silenced. Further work is required to definitively answer this.

Another explanation is that GmYSL7 plays a more direct role in metal ion homeostasis, in line with the role proposed by Castro-Rodríguez et al. (2020, accompanying manuscript). If GmYSL7 transports a peptide that signals cellular metal concentrations, then when transport of the peptide ceases the plant may not be able to sense the metal status of the symbiosome, resulting in an iron starvation response in which transcription factors, among them WRKY9, are expressed and upregulate metal transporters, increasing iron transport to the nodules. If simultaneously symbiosome development stalls, these metals may accumulate outside the symbiosome, requiring storage in other forms, such as ferritin. Further study is required to validate these proposals, including the identification of the peptide substrate for GmYSL7.

## Conclusion

We have identified a member of the oligopeptide transporter family, GmYSL7, which is localized to the symbiosome membrane in nitrogen-fixing soybean nodules. It transports an array of small oligopeptides out of the symbiosome and into the plant cell cytosol, and its disruption arrests infected cell development and symbiosome maturation, inhibiting nitrogen fixation. It affects expression of a number of genes involved in plant defense responses and in iron homeostasis. It is also able to rescue a *M. truncatula* mutant, in which the equivalent gene is compromised, indicating conserved function across the two legumes.

## METHODS

### Plant Growth Conditions

*Glycine max* L. cv Stevens (soybean) seeds were inoculated at planting and one week after planting with *Bradyrhizobium diazoefficiens* (Soybean group H, New Edge Microbials). Plants were grown as described in Clarke et al. (2015) and fertilized once a week with a nitrogen-free B&D nutrient solution (Broughton and Dilworth 1971). Nitrogenase activity in nodules was assessed using an acetylene reduction assay as described by Unkovich et al. (2008).

For limited and excess iron conditions plants were grown in B&D solution with 0, 1, 10 (control concentration) or 100 μM Fe-citrate, which was renewed every 2 days to maintain pH and stable nutrient supply. Two biological replicates were done. Fe status was determined by elemental analysis (Lee M, School of Land and Environment, University of Melbourne) using the Perchloric Nitric Acid Method. 15 plants per treatment were analyzed to determine shoot iron content using an Inductively Coupled Plasma Optical Emission Spectrometer (Varian Medical Systems, Palo Alto, CA, USA).

### Cloning and Constructs

Genomic DNA for cloning the *GmYSL7* promoter was extracted from mature soybean leaves using DNeasy Plant minikit (Qiagen). RNA was extracted from plant tissues using an RNeasy Plant mini kit (Qiagen) and cDNA synthesized using an iScript cDNA synthesis kit (Invitrogen). All constructs were PCR amplified from soybean nodule cDNA or gDNA using either Platinum Pfx50 (Invitrogen) or Phusion (Thermo Fisher Scientific) high fidelity polymerases and cloned using the Gateway cloning system (Invitrogen). A list of primers used can be found in Supplementary Table 3.

For GmYSL7 promoter GUS fusion constructs, a 2 kb genomic fragment immediately upstream of the GmYSL7 coding region was recombined into either pKGW-GGRR (Gavrin et al. 2016) or pKGWFS7 (Karimi et al., 2002). The full-length coding sequence of GmYSL7 was recombined into pGmLBC3-pK7GWIWG2 Gateway vector (Gavrin et al. 2016) to create a hairpin RNAi vector for silencing the gene. N-terminal GFP fusion constructs for GmYSL7 were constructed from the full-length coding sequence recombined into either pGmLBC3-pK7WGF2-R (Gavrin et al. 2016) or a modified pK7WGF2 (pGmLBC3-pK7WGF2) where the 35S promoter is replaced by the GmLBC3 promoter. The free GFP construct was made by *Eco*RV digestion and re-ligation of the pGmLBC3-pK7WGF2 vector to remove the intervening Gateway cassette. For the symbiosome space GFP construct, MtNOD25 (Hohnjec et al. 2009) was PCR amplified from *M. truncatula* cDNA and recombined with pGmLBC3-pK7WGF2. For yeast expression, full length open reading frames of GmYSL7, AtYSL7, AtIRT1, AtOPT4 and ZmYS1 inserted into the pDR196GW vector.

The GmYSL7 coding sequence was synthesised with the MtYSL7 promoter and flanked by attL recombination sites inserted in the pUC57 (Synbio). The construct was recombined into pGBW13 using Gateway Cloning technology.

### Transformation of Soybean and Medicago

Hairy root transformation of soybeans (cv Stevens) used *Agrobacterium rhizogenes* K599 and was as described by Mohammadi-Dehcheshmeh et al. (2014). Transformed roots were inoculated with *B. diazoefficiens* CB1809 (Becker Underwood, Somersby, NSW, Australia). Plants were grown under controlled temperature and lighting conditions (26°C day, 24°C night; 16 hr day; 120-150 μmol m^−2^ s^−1^). Transformed nodules were examined 2-4 weeks post inoculation.

Transformation of *Medicago truncatula* was as described by Boisson-Dernier et al. (2001) using *A. rhizogenes* ARqua1.

### Microscopy

Confocal imaging of GFP-fusion proteins was done on transgenic nodules either hand sectioned or sectioned in low melt agarose using a vibratome (752M Vibroslice, Campden Instruments, Loughborough, Leics., UK). In some instances, nodules were counterstained by FM4-64 (30 μg/ml). Nodule sections were immediately imaged as described previously (Limpens et al., 2009) using either an LSM Pascal 410 (Zeiss) or an SP5 II (Leica) confocal laser-scanning microscope.

Imaging of GUS expression was done as described in Clarke et al. (2015). Sections were either counterstained with ruthenium red or mounted directly in Milli-Q water, and imaged using an Axiophot epifluorescence microscope with a set of Achroplan objective lens (Zeiss).

The protocol for tissue preparation for light and EM has been described previously (Limpens et al., 2009). Semithin sections (0.6 μm) for light microscopy and thin sections (60 nm) for EM of transgenic nodules were cut using a Leica Ultracut ultramicrotome UC7 (Leica). Sections were collected on 400 mesh nickel grids and examined using a Jeol JEM 1400 transmission electron microscope (Jeol Ltd, Tokyo, Japan).

### Quantitative Reverse Transcription-PCR

RT-qPCR assays were used to measure transcript abundance in soybean tissues of control and YSL7 RNAi plants grown in sand or hydroponics. cDNA was synthesized from 500 ng total RNA using Iscript reverse transcriptase (Bio-Rad, Hercules, CA, USA), according to manufacturer’s instructions. Quantitative real time PCR assays were done in a volume of 5 μl in triplicate and contained 1 μl of cDNA diluted 1/5, 1 X LightCycler® 480 SYBR green I mix (Roche Applied Science, Castle Hill, Australia) and 0.5 μM of each primer (GmYSL7 and GmUBI3 QRT primers; Supplementary Table 3). Assays were done using a LightCycler® 480 (Roche Applied Science) and the following conditions: 95°C 10 min, 45 cycles of 95°C 10 s, 56°C 10 s, 72°C 20 s, followed by ramping the temperature from 55°C to 95°C for melt curve analysis. PCR efficiency for each primer pair was determined using the LinRegPCR software (Ramakers et al., 2003) and data analysed using the LightCycler® 480 software package (Roche Applied Science). Data were normalized using *GmUBI3* (Glyma20g27950; Trevaskis *et al*., 2002) or *cons6* expression (Libault et al., 2008). Stable *GmUBI3* expression in the tissues examined in this study was confirmed through comparison of its expression with five characterized soybean reference genes (*cons4*, *6*, *7*, and *15*; Libault et al., 2008) using geNorm software (Vandesompele et al., 2002). The amplified product from the real-time reaction was cloned and sequenced to confirm the specificity of the amplification product.

### Yeast complementation

To test for transport of Fe(II)NA AtYSL7, GmYSL7, ZmYS1 in pDR196GW and the empty vector were introduced into the yeast *fet3/fet4/ftr1* mutant (Spizzo et al. 1997; DEY1530: *MATa ade2 his3 leu2 lys2 trp1 ura3 fet3-2∷HIS3 fet4-1∷LEU2 ftr1D1∷TRP1*) using the method described by Dohmen et al. (1991). Fe(II)-NA plates were prepared by mixing 15 μl 10 mM FeSO_4_ in 200 mM MES/Tris pH 7.4 with 250 μl 200 mM Na-ascorbate and 8 μl of 50 mM NA and heating at 65^°^C for 10 minutes to produce a clear solution that was added to 25 ml of SD media to produce the Fe(II)-NA plate. Transformants were grown in liquid media to an OD600 of 1 and then serially spotted in ten-fold dilutions on either SD plates with no added iron, Fe(II)-NA plates or SD-plates with 10 μM Fe-citrate.

For the peptide transport assay AtOPT4, AtYSL7, GmYSL7 in pDR196GW and the empty vector were introduced into the yeast *opt1* mutant (Y11213: BY4742; *MATα; ura3Δ0; leu2Δ0; his3Δ1; lys2Δ0; YJL212c∷kanMX4*, Euroscarf). Transformants were grown as above on minimal medium (0.17% YNB without amino acids and (NH_4_)_2_SO_4_, supplemented with amino acids as required, and containing either 10 mM NH_4_Cl (positive control) or 100 μM of the following peptides, ALAL, LSKL, IIGLM, KLLLLG, DRVYIHPF, DRVYIHPFHL or RLAPEGPDPHHN, as the sole source of nitrogen.

### Syringolin A transport assay

An assay for transport of Syringolin A by AtYSL7 and GmYSL7 in the yeast strain Δ*pdr5* (Y12409: BY4742; *MATα; ura3Δ0; leu2Δ0; his3Δ1; lys2Δ0; YOR153w∷kanMX4*, Euroscarf) was done as described in Hofstetter et al. (2013). Syringolin A was kindly provided by Robert Dudler, University of Zurich.

### Statistical Analyses

A one-way ANOVA with Tukey’s HSD (SAS Enterprise Guide Version 4.3; SAS Institute Inc., Cary, NC, USA) was used to analyze differences in plant organ dry mass after growth in varying Fe concentrations. Differences are reported as significant where *p* < 0.05.

### RNAseq analysis of transcriptome in GmYSL7-RNAi nodules

The transcriptome for *GmYSL7*-RNAi nodules was compared to those transformed with an empty vector control using RNAseq. Hairy root transformation with pGmLBC3-pK7GWIWG2 vector or the vector containing *GmYSL7* coding sequence was used to produce transformed nodules. RNA was isolated from nodules 21 or 22 days after inoculation using an RNeasy kit (Qiagen). Five replicates for each construct were done, each with nodules from 5-7 transformed plants. RNA integrity number (RIN) was determined on a 2100 bioanalyzer (Agilent) and was between 7 and 8.3 for all samples. RNAseq library construction and analysis were completed at Institute for Molecular Bioscience Sequencing Facility, The University of Queensland. A combination of the Ribo Zero rRNA removal bacteria (Illumina) and Ribo Zero rRNA removal plant (seed and root) (Illumina) was used to eliminate the rRNA from the sample. The library was constructed using a TruSeq® Stranded mRNA LT - SetA and SetB (Illumina). Sequencing was performed using the Illumina NextSeq500 (NextSeq control software v1.4/ Real Time Analysis v2.1). The library pool was diluted and denatured according to the standard NextSeq protocol, and sequenced to generate single-end 76 bp reads using a 75 cycle NextSeq500/550 High Output reagent Kit (Illumina).

Raw sequence reads were aligned to the JGI Wm82.a2 soybean assembly. DESeq2 (Love et al. 2014) was used to test for differential expression between control and YSL7-RNAi samples. Genes with log_2_ fold change (log_2_ FC) >1 and adjusted p-value < 0.05 were considered differentially expressed. Overrepresented biological terms were identified from the list of differentially expressed genes. GO term enrichment analysis was based on the information in SoyBase (https://soybase.org/). Enriched biological terms and their linkage were analysed and visualized using ClueGO v2.5.5 (Bindea et al., 2009), implemented in the Cytoscape v3.5.1 environment (Shannon et al., 2003; https://cytoscape.org/cy3.html). ClueGO parameters were as follow: Analysis Mode, Functional Analysis; Load Markers List, Glycine max (3847); Visual Style: Significance Shape, ellipse; ClueGO settings, Ontology/Pathway; GO, Biological Process / KEGG: downloaded the 12/12/2017; Evidence type, All Evidences; Statistical Test Used = Enrichment/Depletion (Two-sided hypergeometric test); Correction Method Used = Bonferroni step down; Min GO Level = 3, Max GO Level = 8, Min Percentage = 4.0, GO Fusion = true, GO Group = true, Kappa Score Threshold = 0.4; Over View Term = SmallestPValue; Group By Kappa Statistics = true; Initial Group Size = 1;Sharing Group Percentage = 50.0.

### Symbiosome isolation, SM and microsomal membrane isolation and proteomic analysis

Symbiosomes were isolated as described by Clarke et al. (2015). SM was collected after pelleting of the membrane and resuspended in 1M Urea for proteomic analysis. Microsomal membrane was isolated from nodules ground and filtered through miracloth as described in Clarke et al. (2015). Symbiosomes and other intact organelles were pelleted by centrifugation at 20000 g. The membrane in the supernatant (enriched in PM and endoplasmic reticulum) was collected by centrifugation at 100,000 g for 1 hour at 4°C and the pellet resuspended in 8M Urea. Proteomic analysis was completed at the La Trobe Comprehensive Proteomics Platform (La Trobe University). Data were collected on a Q Exactive HF (Thermo-Fisher Scientific) in Data Dependent Acquisition mode using *m*/*z* 350–1500 as MS scan range at 60 000 resolution. HCD MS/MS spectra were collected for the 7 most intense ions per MS scan at 60 000 resolution with a normalized collision energy of 28% and an isolation window of 1.4 *m*/*z*. Dynamic exclusion parameters were set as follows: exclude isotope on, duration 30 s and peptide match preferred. Other instrument parameters for the Orbitrap were MS maximum injection time 30 ms with AGC target 3 × 10^6^, MSMS for a maximum injection time of 110 ms with AGT target of 1 × 10^5^.

Raw files consisting of high-resolution MS/MS spectra were processed with MaxQuant version 1.5.5.1 to detect features and identify proteins using the search engine Andromeda. Sequence data for soybean from Phytozome (https://phytozome.jgi.doe.gov/pz/portal.html#!info?alias=Org_Gmax) was used as the database for the search engine.

## ACCESSION NUMBERS

The accession number for *GmYSL7* is NM_001289202.2

## ACKNOWLEGMENTS

We thank Catherine Curie for providing the plasmid containing ZmYS1 and useful discussions about YSL transporters and Sarah Conte and Elsbeth Walker for providing advice about the methods for yeast assays for transport of Fe(II)-NA.

## SUPPLEMENTAL MATERIAL

Supplementary table S1. Unique GmYSL7 and GmNOD26 peptides^1^ identified in purified symbiosome membrane, a microsomal membrane fraction, and a symbiosome-enriched membrane sample from soybean nodule homogenate.

Supplementary table S2. Genes upregulated or downregulated in GmYSL7-RNAi nodules and data for all genes expressed in the nodules.

Supplementary table S3. Primers used in this study. Supplementary figure S1. Phylogenetic analysis of YSL proteins

Supplementary figure S2. Expression analysis of the YSL genes in soybean.

Supplementary figure S3. Expression of *GmYSL7* in response to Fe status during nodule development.

Supplementary figure S4. Detailed morphological analysis of determinate nodule development

Supplementary figure S5. FM 4-64 stained symbiosomes extracted from empty vector control and GmYSL7-RNAi nodules.

## Notes

**Funding information :** This research was funded by the Australian Research Council Discovery Projects DP0772452, DP120102780 and DP150102264 and Industrial Transformation Research HUB IH140100013.

